# Genetic variation/evolution and differential host responses resulting from in-patient adaptation of *Mycobacterium avium*

**DOI:** 10.1101/295121

**Authors:** N. Kannan, Y.-P. Lai, M. Haug, M. K. Lilleness, S. S. Bakke, A. Marstad, H. Hov, T. Naustdal, J. E. Afset, T. R. Ioerger, T. H. Flo, M. Steigedal

## Abstract

*Mycobacterium avium* (Mav) complex (MAC) are characterized as non-tuberculosis mycobacteria and are pathogenic mainly in immunocompromised individuals. MAC strains show a wide genetic variability, and there is growing evidence suggesting that genetic differences may contribute to a varied immune response that may impact on the infection outcome. The current study aimed to characterize the genomic changes within Mav isolates collected from single patients over time and test the host immune responses to these clinical isolates. Pulsed field gel electrophoresis and whole genome sequencing was performed on 40 MAC isolates isolated from 15 patients at the Department of Medical Microbiology at St. Olavs Hospital in Trondheim, Norway. Patients (4, 9 and 13) who contributed more than two isolates were selected for further analysis. These isolates exhibited extensive sequence variation in the form of single nucleotide polymorphisms (SNPs), suggesting that Mav accumulates mutations at high rates during persistent infections. Infection of murine macrophages and mice with sequential isolates from patients showed a tendency towards increased persistence and down-regulation of inflammatory cytokines by host-adapted Mav strains. The study revealed rapid genetic evolution of Mav in chronically infected patients accompanied with change in virulence properties of the sequential mycobacterial isolates.

**IMPORTANCE:** MAC are a group of opportunistic pathogens, consisting of Mav and *M. intracellulare* species. Mav is found ubiquitously in the environment. In Mav infected individuals, Mav has been known to persist for long periods of time, and anti-mycobacterial drugs are unable to effectively clear the infection. The continued presence of the bacteria, could be attributed to either a single persistent strain or reinfection with the same or different strain. We examined sequential isolates collected over time from Mav infected individuals and observed that most patients carried the same strain overtime and were not re infected. We observed high rates of mutation within the serial isolates, accompanied with changes in virulence properties. In the light of increase in incidence of MAC related infections, this study highlights the possibility that host adapted Mav undergo genetic modifications to cope with the host environment and thereby persisting longer.

---

*Mycobacterium avium* (Mav) complex (MAC) are a group of opportunistic pathogens, consisting of Mav and *M intracellulare* species, which primarily affect individuals with weakened immune systems (1–4). However, even healthy individuals can be infected by Mav and in healthy children the disease manifests as lymphadenitis (5–7). Mav infections are very hard to treat, and many anti-mycobacterial drugs fail to clear the infection even after prolonged treatment of 18-24 months (8, 9).

The host response to *Mycobacterium tuberculosis* (Mtb) has been extensively examined over the years. Infections with non-tuberculous mycobacteria are less well characterized, although the focus has been increasing in recent years (10). Mav lacks several of the key virulence factors of Mtb, but can still establish chronic infections. Macrophages are central in defense against mycobacterial infections but they also end up hosting the pathogens as these manipulate cell-autonomous host defenses. When infected, macrophages release interleukin (IL)-12 that aids in activation of T cells, leading to interferon gamma (IFNγ)-producing CD4+ T helper 1 (Th1) cells, thought to be essential in fighting mycobacterial infections (11). IFNγ acts back on the macrophages and strengthens their antimicrobial capacities. Other factors such as tumor necrosis factor alpha (TNFα) produced by activated macrophages and T cells, are not only important in enhancing the microbicidal capacity but also in inducing the adaptive response, in synergy with IFNγ (12, 13). Pro-inflammatory cytokines like IL-6 and IL-1β have also been shown to play a role in mycobacterial infections (11, 14–17). IL-1β along with IL-6 and TNFα have been observed to be suppressed by the most virulent strains of Mav (18).

The genome of Mtb was thought to be relatively unaffected by the host environment (19), but recent data from macaques suggest that Mtb acquires mutations during long term Mtb infection (20). Genomic analysis has revealed greater genomic heterogeneity in Mav than can be seen in Mtb (21–23). This is probably a consequence of the variety of niches that Mav species occupy (such as soil, fresh water, showerheads etc.), and also indicates that Mav may be more prone to variability than Mtb. Genetic change within an infected patient has not been investigated for Mav infection. The effect of genetic variation on virulence and pathogenesis is even less understood. An early study on Mav virulence found that strains varied in virulence depending upon where they were isolated from, despite having very similar genomic composition (24, 25). Genetic changes could be selected for when they provide increased virulence and persistence. We hypothesized that the hostile environment during chronic Mav infections in humans would result in genetic changes of the infecting Mav strain, and possibly alter the virulence of the strain. In the present study, we investigated genetic changes by sequencing Mav isolates sampled from individual patients over time, and studied host responses to these sequential isolates *in vitro* in primary mouse macrophages and *in vivo* in mice.

## RESULTS

### Sequential sampling from patients with MAC infection shows persistent infection over time

For this study, 15 patients diagnosed with MAC disease were monitored over a period of one month to 3 years. At least two consecutive bacterial samples were isolated from all patients with at least one month interval, providing a total of 40 isolates (Table S1). To differentiate between the two MAC species (Mav and *M. intracellulare*), melting curve analyses of 16S rRNA from the 40 isolates were compared to type strains of Mtb, Mav and *M. intracellulare* (Fig. S1). Thirty one isolates were identified as Mav and nine as *M. intracellulare*. All patients had the same species identified in all isolates, suggesting either the same bacterium persists over time or reinfection with the same species. To evaluate whether strains isolated from a patient at different times represented persistent infection or reinfection with a new strain of the same species, we performed PFGE using SnaBI restriction analysis (Fig1a). Sequential isolates were found to be of indistinguishable genotypes in 13 of the 15 patients, suggesting persistent infection by a single bacterial strain. Phylogenetic cluster analyses based on PFGE profiles were carried out for each species separately (Fig. 1a). All Mav and *M. intracellulare* isolates showed distinct and similar SnaBI profiles. For the patients 4, 9 and 13, where there are more than two consecutive Mav isolates from the same patient, we observed that most of the isolates have identical (patient 4 and 9) or very similar (Patient 13) SnaBI profiles, suggesting the presence of a single strain persisting over time. For patient 13, the presence of an extra band in the DNA profile of isolates 13.3 and 13.4 indicates that a genetic change occurred sometime between sampling of isolates 13.2 and 13.3.

**FIG. 1.**
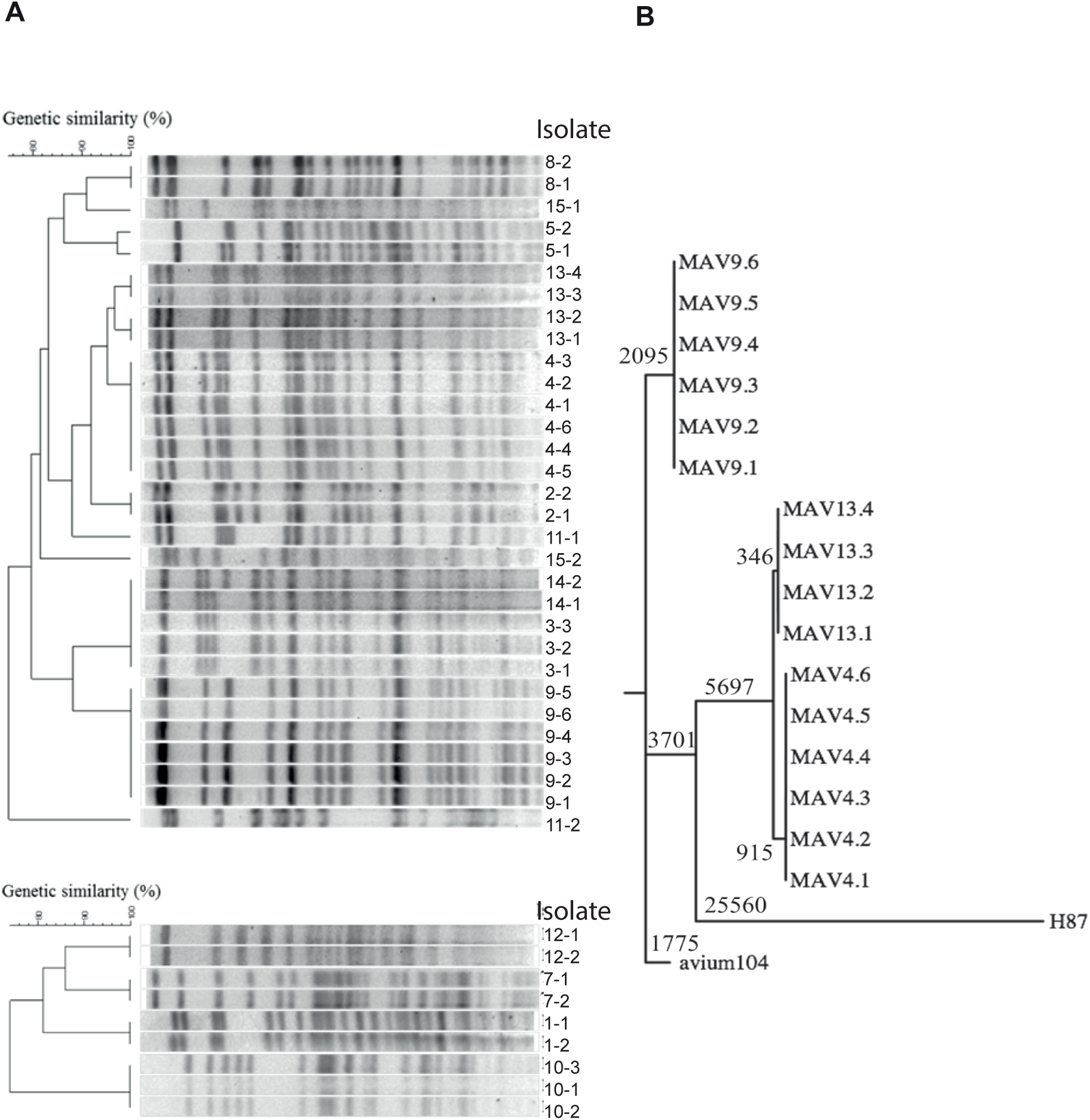
Genetic comparison / classification of clinical MAC isolates from patients. (A) Pulse field gel electrophoresis of 40 clinical isolates from 15 patients was performed using a SnaBI typing method. Restriction enzymatic digestion by SnaBI created distinct profiles that could be used to distinguish between isolates from different patients and between isolates taken at different time points from the same patient. Dendrograms were generated based on cluster analysis using the UPGMA method and Dice similarity coefficient, to assist in visualizing SnaBI pattern similarity.(B) Maximum parsimony tree showing the phylogenetic relationship among the clinical isolates and two Mav reference strains, 104 and H87. The branch lengths indicate the number of changes (SNPs).

### Whole-genome sequencing reveals accumulation of SNPs and genetic variation in Mav during persistent infection

Genetic changes in some of the serial intra-patient isolates were observed from PFGE analysis. To further investigate genetic changes, whole-genome sequencing (WGS) of the isolates was performed. The genome sequences were analysed for polymorphisms like SNPs, using Mav 104 as a reference strain, and a phylogeny tree was constructed. We observed that the isolates from patient 9 were closely related to Mav 104, whereas isolates from patient 4 and 13 were closely related to Mav strain subsp. *hominissuis* H87 (26) (Fig. 1b). Overall, the genome sequences of intra-patient isolates were highly similar, with only 0-17 SNPs between any intra-patient pair, strongly supporting that each patient maintained a unique infecting strain (Fig. 2).

**FIG. 2.**
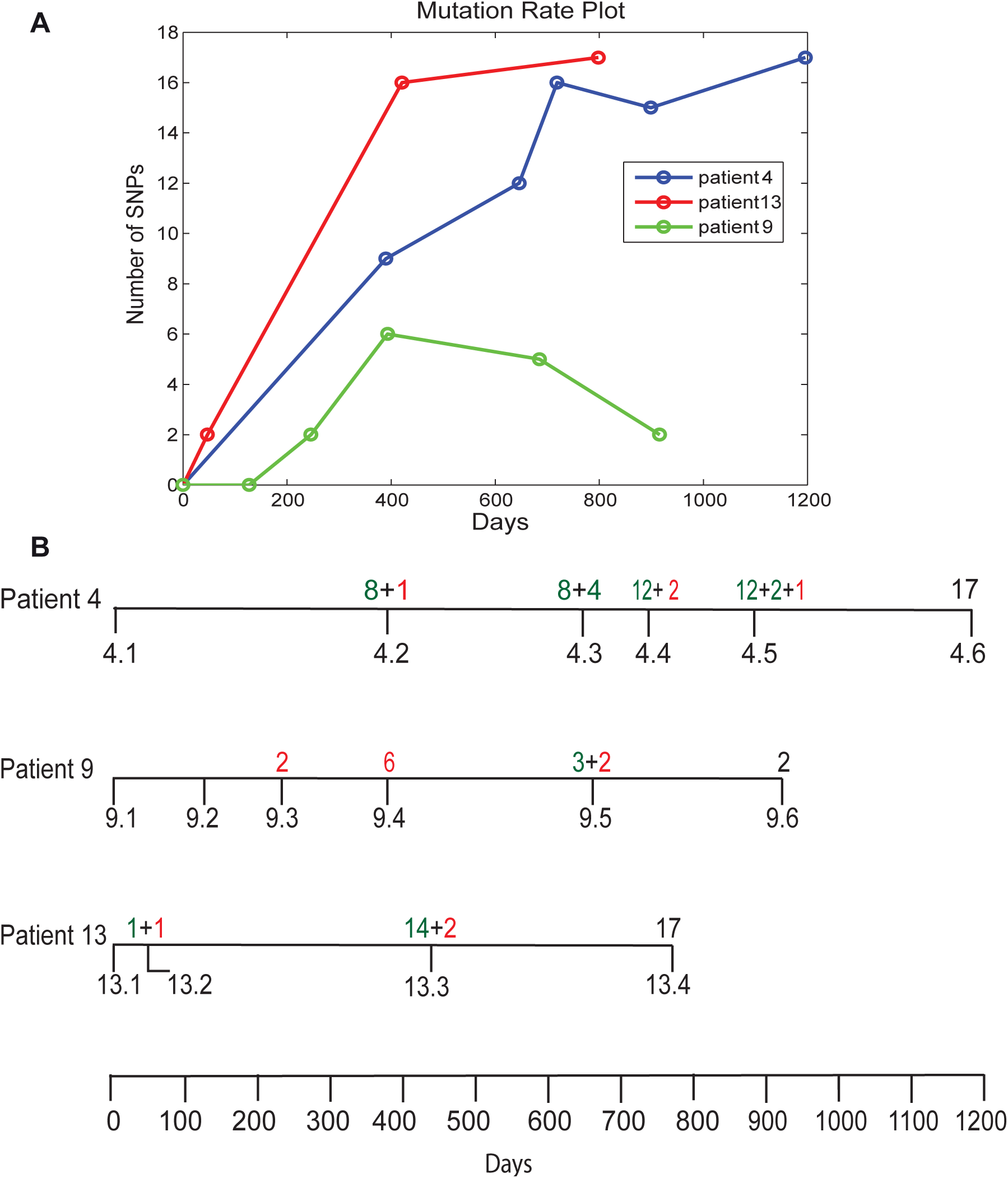
SNPs from whole-genome sequencing of Mav clinical isolates sampled over time from single patients. (A) The maximum-likelihood estimate of the rate parameter for clinical isolates from patients 4, 9 and 13 was calculated, assuming a Poisson model for the accumulation of SNPs over time, and represented as a mutation rate plot using Matlab. (B) Diagrammatic representation of the time-line of sample collection from patients 4, 9 and 13 and corresponding SNP development. Number of SNPs in green represents fixed mutations, whereas numbers in red are unique to that particular isolate and are lost in the subsequent isolates.

To establish the genetic variation between intra-patient isolates of Mav, we considered patients where more than two consecutive isolates were collected. Patient 4 and 9 provided six isolates, whereas patient 13 provided four isolates. The mutation rate in intra-patient isolates was determined by calculating the maximum-likelihood estimate (ML) for the accumulation of SNPs over time. The ML estimates of the rates were 6.92 (95% confidence interval (CI): 4.81-9.64), 2.40 (95% CI: 1.24-4.19) and 12.39 SNPs per genome per year (95% CI: 8.73-17.07) for patient 4, 9 and 13 respectively. This results in an estimated mutation rate of 1.41×10^−6^, 0.44×10^−6^ and 2.41×10^−6^ SNPs per site per year for patients 4, 9 and 13 (Fig. 2a). These rates are higher than mutation rates among clinical isolates from related species, such as Mtb (~ 0.5 SNPs per genome per year) (27). For Mav infections, mutation rates within a single infected patient have not been studied previously.

In patient 4, a comparison of later isolates to the first isolate, 4.1, revealed an accumulation of 17 SNPs over a period of 1196 days. In addition, we observed that 13 of the SNPs were maintained in subsequent isolates, indicating a high degree of fixation of the mutations in the infecting strain (Fig. 2b). The accumulation of SNPs in isolates from patient 4 steadily increased with time, with 9 SNPs observed at the second time-point (4.2, 390 days), 8 of these SNPs were maintained in all successive time points analysed. When 13.1 was used as a reference for isolates from patient 13, the subsequent isolates exhibited a similar accumulation of SNPs as in patient 4, with 17 SNPs in total detected within 4 isolates over a period of 798 days (Fig. 2b). The majority of SNPs occurred over the first year between isolate 13.1 and isolate 13.3, where 14 SNPs occurred that were conserved till the last isolate 13.4. For patient 9, the pattern of SNPs in consecutive isolates was different. Although a total of 13 SNPs were identified for the series of isolates, only 2 mutations were fixed in subsequent isolates. We observed no differences in preservation between synonymous and non-synonymous mutations that could explain the difference in the level of fixation in subsequent isolates (Table S2).

Since we observed an additional band in the PFGE for patient 13 (Fig 1a), we compared all the isolates from patient 13 including 13.1 to Mav 104 as an outgroup and observed that some SNPs occurring in 13.3 and 13.4 were associated with 13.1 and 13.2. In addition, sequencing revealed that a putative prophage (MAV0799-MAV0845) appears to be deleted in 13.1 and 13.2, but not 13.3 and 13.4. These observations together indicate the presence of 2 subtypes that may have originated within patient 13. Due to lack of multiple samples at each time point, it is difficult to further substantiate this observation. Similar analysis was performed for patients 4 and 9. For patient 9, the SNPs did not change with respect to Mav 104, however for patient 4, 4 mutations now appeared to associate with 4.1 instead of 4.2-4.6 (Table S3). This suggests that there are 2 subtypes in patient 4 as well. The re-calculated mutation rates based on the assumption of sub lineages for patients 4 and 13, resulted in 6.6 and 7.1 SNPs per genome per year, respectively (Fig S5).

### Host-adaptation can change the ability of Mav to survive in murine bone marrow derived macrophages

In order to evaluate whether changes within the isolates from patients 4, 9 and 13 were a response to host adaptation, we first studied the ability of these bacteria to grow in culture (Fig. 3a). Only minor differences in growth rates were observed for isolates from patient 4 and 9. However, for patient 13 we observed relative growth impairment for the first isolate when compared to the other 3 isolates (Fig. 3a). Isolates were next compared for intracellular growth in murine macrophages over 7 days. Enumeration of bacteria (colony forming units, CFU) 2 hours post infection (p.i) suggested equal uptake for all isolates (Fig. S2). For patient 9 and 13, a significant increase in CFU between the first and the last isolate was observed (Fig. 3b-c). For patient 9, CFU at day 7 increased 1.7 fold from 9.7×10^6^ for 9.1 to 1.6×10^7^ CFU for 9.6, and for patient 13 CFU increased 7.8 fold from 8.5×10^6^ for 13.1 to 6.6×10^7^ for 13.4 from day 3 to day 7 p.i, suggesting that these strains increased their virulence during infection. For patient 4, no difference was observed in the ability to survive inside macrophages (2.6×10^6^ CFUs for 4.1 vs. 3.0×10^6^ CFUs for 4.6). Taken together our results suggest that Mav dynamically adapts to the hostile environment of the host, thus facilitating persistence /chronic infection.

**FIG. 3.**
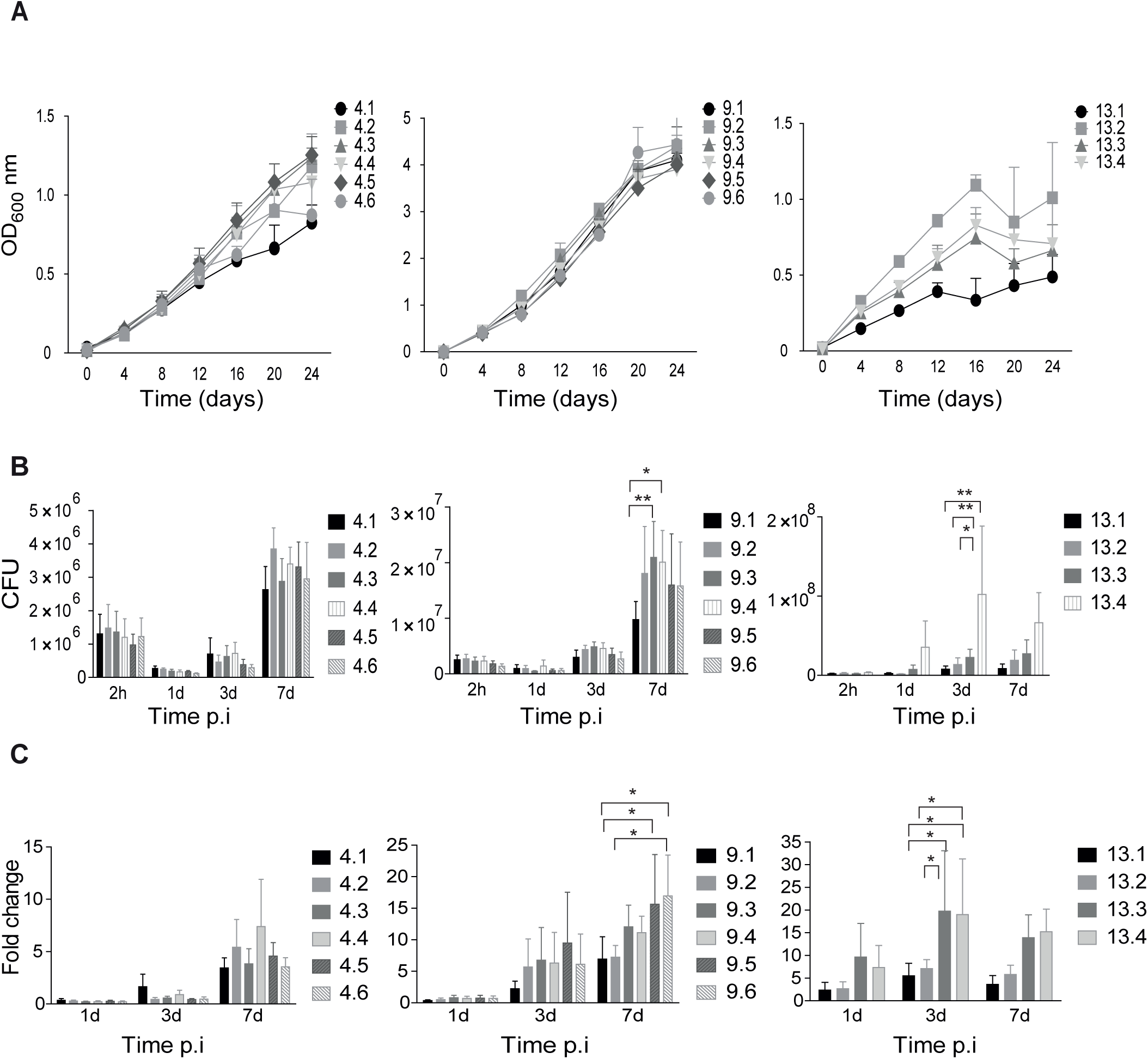
Growth in broth and intracellular replication in macrophages of sequential Mav isolates A. Growth curves were recorded of sequential Mav isolates from patients 4, 9 and 13 grown for a period of 24 days. Isolates are numbered chronologically from the time they were collected from the patients. Recordings were performed in triplicates; the OD values show mean ± SEM. B. Intracellular replication in murine BMDMs infected with isolates from patients 4, 9 and 13 at a MOI of 10. CFU counts from lysed macrophages were determined 1, 3 and 7 days after infection. (C) Intracellular bacterial counts represented as fold change normalized to the uptake of bacteria, at 2 hours post exposure /infection. Bars represent mean ± SEM from 4 independent experiments; * p<0.05, **p<0.01 by Repeated measures Two-way ANOVA, Tukey post-test.

### Host-adaptation can change the inflammatory properties of Mav

Inflammatory responses are central in controlling infection and we have previously shown that negative regulation of inflammatory responses by Keap1 or by depletion of Toll-like receptor (TLR)-signaling facilitates intracellular replication of Mav in human primary macrophages (28–30). We therefore performed a broad systematic screen on the early expression of inflammatory genes to identify changes in immune stimulatory properties that could explain the change in survival. Murine BMDMs were infected with the first and the last clinical isolates of patient 9 at a MOI of 10 for 6 hours. Genes were identified and plotted that were differentially expressed (at least two-fold induced or repressed) in macrophages infected with the last compared to the first isolate (Fig.S3). The analysis revealed down-regulation of many pro-inflammatory cytokines and we decided to measure protein levels of two of them, IL-6 and IL-1β, from supernatants of BMDMs infected with all isolates from patient 4, 9 and 13 (Fig. 4a-b). IL-1β secretion was low and highly variable for all strains, whereas IL-6 secretion was more potently induced. As the gene expression data indicated, we observe that despite an increased bacterial load in the macrophages infected with the final isolates of patient 9 and 13, the IL-6 levels were lower than from supernatants of macrophages infected with the first isolates. These results are in line with previous studies (31), and the reduced inflammation may facilitate intracellular survival (Fig. 3). However, patient 4 isolates did not show a similar reduction in cytokine production and confirmed the level of variation that has been previously reported (29).

**FIG. 4.**
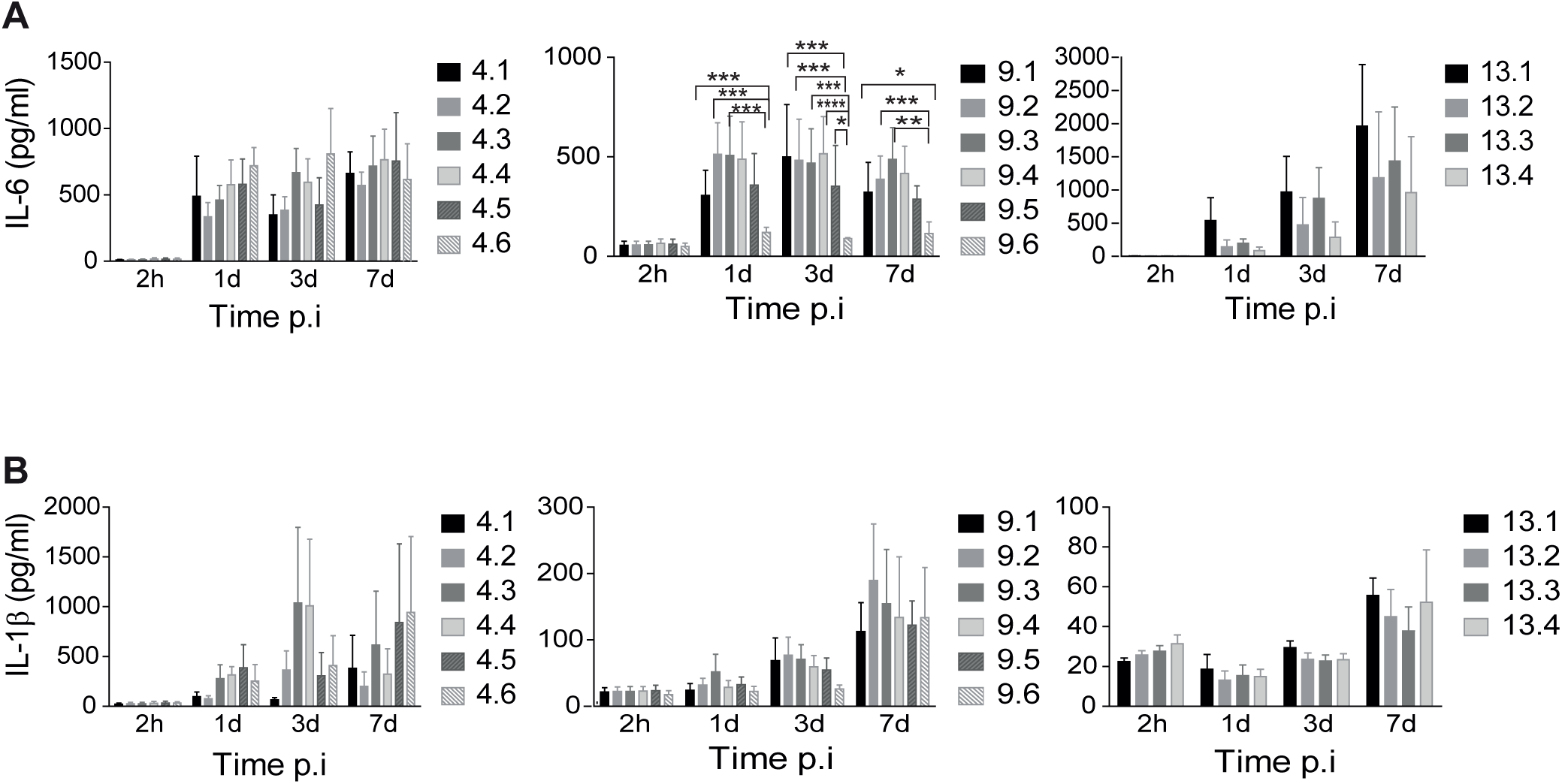
Down-regulation of pro**-**inflammatory cytokines in Mav infected mouse macrophages. Murine BMMs were infected with the sequential Mav isolates from patient 4, 9 and 13 at a MOI of 10. Levels of IL-6 (A) and IL-1β (B) were measured from supernatants at 2 h, 1 day, 3 days and 7 days post infection (p.i). Bars represent mean ± SEM from three or four independent experiments; *p<0.05, **p<0.001, ***p<0.0001 by Repeated measures Two-way ANOVA, Tukey post-test.

### Host-adaptation can change the ability of Mav to survive in mice

To validate if *in vitro* observations were reflected *in vivo*, we infected C57BL/6 mice with isolates from patient 9 (9.1, 9.5, 9.6) and 13 (13.1, 13.2, 13.4) (Fig. 5). We chose to investigate isolates from patient 9 and 13, as the *in vitro* data strongly indicated that the last isolates from these patients showed increased survival within the macrophages compared to the early isolates. At day 15, 30 and 70 of infection, we measured the number of bacteria from both spleen and liver of mice (Fig. 5a and b respectively). Overall, the bacterial loads were constant or slightly increasing over time for most isolates in both liver and spleen.

**FIG. 5.**
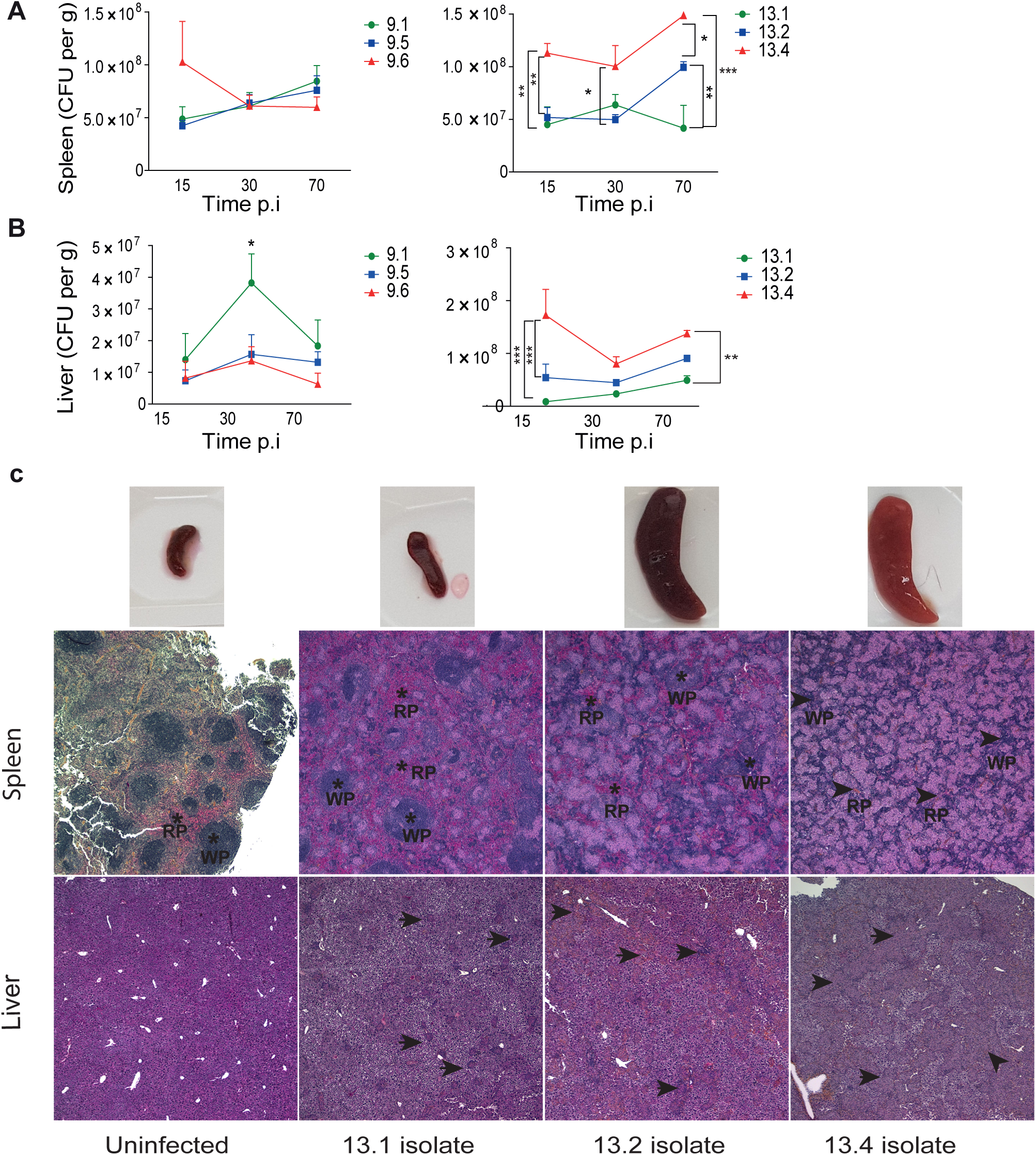
Mycobacterial load and tissue pathology in mice infected with sequential Mav clinical isolates. (A-B) C57BL/6 mice were infected intraperitoneally with Mav isolates from patient 9 (9.1, 9.5, 9.6) and patient 13 (13.1, 13.2, 13.4) for 15, 30 and 70 days. Tissue bacterial loads in spleen and liver are shown as mean CFUs per gram ± SEM from two experiments with four mice in each group. *p<0.05, **<0.01, ***<0.001 by Repeated Measures Two-way ANOVA and Bonferroni’s post-test. (C) Histological examination of spleen and liver from mice infected with Mav isolates 13.1, 13.2 and 13.4 at day 70 post infection. Upper and middle Panel: spleens and spleen tissue histology sections. RP indicates the red pulp and WP the white pulp areas. Lower Panel: liver histology. Arrows point to granulomatous structures. Images are taken at 40x magnification. Experiments done twice with similar results, one representative experiment is shown.

For patient 9, the only significant difference was seen in the liver at day 30 p.i, where there was an increase in CFU for the first isolate when compared to the last isolates (9.5, 9.6) (p<0.05). This is incongruent with our observations in macrophage infections (Fig. 3b-c). However, the *in vivo* data suggests no overall difference in virulence between the three isolates from patient 9(Fig. 5a and 5b, left). For Patient 13, the *in vivo* condition reflected our observations of increased infectivity that we made in macrophages (Fig, 3b,c). Throughout infection, we found significantly increased bacterial loads in liver and spleen from mice infected with the last isolate, 13.4, compared to 13.1 and 13.2 (Fig. 5a and b, right), suggesting a gain in virulence over time. At day 70, spleen size increased substantially in mice infected with isolates 13.2 and 13.4 compared to 13.1, and the color of the 13.4 spleens was strikingly pale when compared to the other spleens (Fig. 5c, upper row).

On further histological examination of spleens on day 70 p.i. (Fig. 5c, middle row), we observed that the structure of the 13.4 spleens was almost replaced by coalescing granulomas. In contrast, smaller amounts of single small or medium sized granulomas, in part coalescing for 13.2 spleens, were present in 13.1 and 13.2 spleens on day 70 p.i. Myeloid precursor cells and megakaryocytes were found in all affected spleens, indicating extra medullary hematopoiesis. A similar pattern of granulomatous infiltration was seen in liver sections on day 70 p.i. (Fig. 5c, lower row), however the distinction between normal tissue and granulomas was less obvious than in the spleen. Infection with the 13.4 isolate resulted in replacement of much of the normal liver tissue with sheets of granulomas. The granulomas were sparse and mostly separated for 13.1, the granulomas were moderate in number for 13.2 but starting to coalesce. It is important to note that though 13.1 showed impaired growth *in vitro*, *in vivo* it still managed to survive, albeit poorly when compared to 13.4. Taken together our data suggest that the Mav isolated from patient 13 changed properties over the course of infection in the patient and increased its ability to survive. Finally we investigated cytokine levels in organ homogenates from infected mice. The decrease in level of IL-6 and IL-1β observed in macrophages could not be observed in liver homogenates where levels were steady over time and between isolates (Fig. S4).

### Splenic CD4+ and CD8+ effector cytokines decrease over time in mice infected by all Mav isolates from patient 9 and 13

It has previously been demonstrated that Mav infections elicit a CD4+ Th1 immune response, which is associated with resisting the spread of the pathogen (32, 33). In our study, we analyzed Mav-specific IFNγ or TNFα (effector cytokine) production from CD4+ and CD8+ T cells that were isolated from spleens of infected mice (Fig. 6). We observed that the frequency of both Mav-specific CD4+ and CD8+ T cells were high on day 15 p.i. and rapidly decreased by day 30 for patient 9, and more gradually to day 70 p.i for patient 13. This decrease in Mav-specific T cells was observed with all tested Mav isolates, despite sustained or even increased tissue bacterial loads (Fig. 5), The frequency of cytokine-producing CD8+ T cells was lower (<5%) than CD4+ T cells (20-40%) and did not change much over the course of infection.

**FIG. 6.**
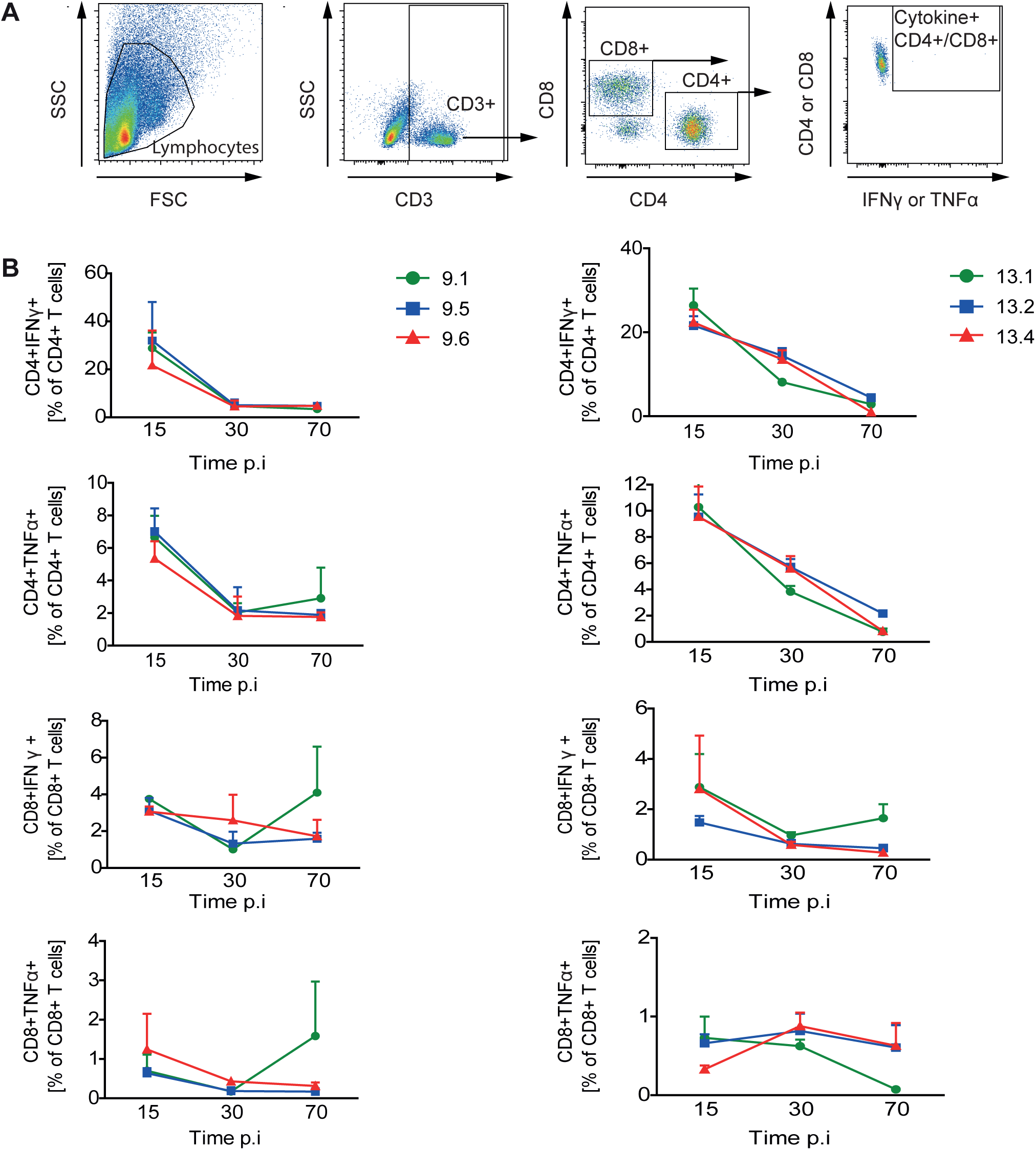
Mav-specific splenic T cell responses. C57BL/6 mice were infected with isolates of patient 9 (9.1, 9.5, and 9.6, left) and patient 13 (13.1, 13.2, and 13.4, right) for 15, 30 and 70 days. Mav-specific T cell responses were measured from *in vitro* Mav re-stimulation of splenocytes 15, 30 and 70 days post infection (p.i). IFNγ and TNFα production from CD4+ and CD8+ T cells were analyzed by intracellular flow-cytometry. A) Gating strategy. B) Results are represented as percentage of T cells producing the indicated cytokine. Results show mean ± SEM of two experiments with four mice analyzed in each group.

## DISCUSSION

In the current study, we evaluated the mutations acquired by Mav during persistent infection in human patients and further tested the pathogenesis of these isolates in macrophages and mice. Within macrophages, the Mav isolates from patients 9 and 13 exhibited higher bacterial load and down-regulated inflammatory cytokines. However, the last isolates from patient 13, but not patient 9, showed increased survival in mice compared to the first isolate.

Most intra-patient isolates showed similar PFGE profiles, suggesting persistent infection with a single MAC strain. On WGS analysis, we found variable mutation rates in sequential intra-patient isolates. If we compare strains from different patients our findings are in congruence with previous studies, which have reported high genetic diversity between Mav isolates (34–36). The mutation rate observed for isolates within a patient, ranged between 2.4-12.3 SNP’s per year, which is comparatively higher than the mutation rate of ~0.5 SNP reported for Mtb isolates from humans (27) and macaques (20, 37). These observations further indicates that the genome of Mav is more malleable than that of Mtb, and more prone to variation, maybe due to influence from host factors. In addition, we analyzed the patient isolates for SNPs in two ways, the first was to use the first isolate sequence from each patient as the founder strain and tabulate the SNPs present in the subsequent isolates (Table S2). For the second analysis, all isolates from patients were treated as independent cultures and compared it to the reference strain Mav 104 as an out group. For patients 4 and 13, in the second analysis, two sub clonal populations were uncovered (Table S3), which could either be due to a microevolution of a single founder strain or mixed/re-infection. The plausibility of the former is more likely than the later, as PFGE patterns are nearly identical and the WGS pattern of differences found within sequential isolates were identical to each other. The likelihood of being re-infected with such a genetically similar strain is low based on the previous finding, describing high degree of genetic diversity in environmental isolates (38). In case of patient 4 and 13, it can be speculated that the evolution of the isolates may be linear or divergent. For patient 9, the predominance of unique SNPs in individual isolates that were not propagated to other isolates over time (i.e. not fixed), argues against linear evolution, and instead supports heterogeneity of the population structure *in vivo*. Studies in Mtb have examined the presence of sub-clones and intra-patient microevolution (39–42) and support the presence of divergent populations, which can be either generated due to antibiotics (40) or clinical severity (42). Thus, our analysis supports two hypotheses, linear and divergent microevolution.

Innate immune cells like macrophages are the first line of defense against mycobacterial infections. In Mav pathogenesis, like Mtb, macrophages are the primary host cell that initialize the containment of the bacterium (14, 28). We tested intracellular growth within murine macrophages and observed that though all isolates managed to survive within the macrophage, the last isolates collected from two patients (9 and 13) survived better than initial isolates from the same patient. This increased survival of certain isolates could either be due to their increased multiplication rate or impaired ability of the macrophages to eradicate the bug (5, 24, 43, 44). In our experiments, we measured reduced levels of pro-inflammatory cytokines like IL-6 and IL-1β when macrophages were infected with later isolates compared to the initial isolate from a patient. The levels of IL-1β were highly variable in our experiments. Efficient IL-1β production requires that mycobacteria come in contact with the cytosol and since Mav is not present in the cytosol (30), this could explain the varying results. The ability to activate host defenses by Mav has previously been shown to vary between Mav isolates, and down-regulation of pro-inflammatory cytokines is believed to promote survival of the bacterium (12, 18, 45).The present study is the first to indicate that the ability to activate host-defenses can change over the course of an infection by Mav. To support our *in vitro* observations, mice were infected with isolates from two of the patients and infection showed the same trend as for the *in vitro* experiments for one set of isolates. It has been previously observed that there is a discordance between *in vitro* and *in vivo* data (5, 24). One explanation could be that *in vitro* macrophage infections represent an isolated system, as opposed to mice, where other immune cells, including the adaptive arm of the immune system, play an active role in curtailing the infection.

Cytokines like IFNγ and TNFα, produced by effector T cells, contribute to control of the infection at chronic stages (32). We previously demonstrated that, in mice infected with Mav 104, the induction of effector T cell responses coincided with a decrease in bacterial loads (33). In our current experiments, all the clinical isolates from both patients 9 and 13 impaired CD4+ effector T cell responses from around day 30 p.i. and exhibited bacterial persistence over time. Although, *in vitro*, there was an inverse relationship between bacterial count and inflammatory response, *in vivo*, the increase in bacterial counts could not be explained by production of inflammatory cytokines. We speculate that there could be other cells or effector molecules that play a role and that have not been studied, such as B cells, γ/δ T cells, neutrophils, natural killer cells and effector molecules like IL-10, IL-17 and TGF-β which all have previously been reported to have an important role in host defenses towards mycobacteria (46–50). Furthermore, the increased virulence exhibited by the isolates could be due to the high bacterial load. Studies have illustrated that at chronic stages of infection, due to prolonged exposure to high doses of antigen, T cells may undergo terminal differentiation and in parallel undergo apoptosis (51, 52). Considering the results obtained, we speculate that a diverse milieu can lead to modifications in the immune-modulating properties and intracellular proliferation of intra-patient isolates over time. Looking at WGS data (Table S2 and S3) for isolates from patient 13, 13.4 exhibited a distinct increase in survival *in vivo* after the first time-point, in which a SNP in MAV0182 (K38T) was observed. MAV0182 is annotated as a super oxide dismutase (Fe-Mn) (SOD); SODs catalyze the dismutation of the superoxide radical to H_2_O_2_ and oxygen. In *M. paratuberculosis* and Mtb, SOD is actively secreted and has shown to generate protective cellular immunity(53, 54). We speculate a similar role for MAV0182 and the SNP may aid in survival of 13.4. In addition, in 13.2, which also exhibited increased growth, a SNP F267V in MAV2838 was observed. In fact, this was the only novel polymorphism observed in 13.2 compared to 13.1. MAV2838 is an orthologue of the *oxyR* transcription factor, which is known to function during oxidative response. (55, 56). It is conceivable that the mutation F267V in MAV2838 confers a growth advantage through an adaptive response to oxidative stress. Further evaluation of the phenotypic effects of these SNPs, such as recombineering, will be required to test their role in survival.

This is the first study analyzing in-patient evolution of Mav infection in humans. A possible limitation is that the cohort studied was small. In addition, only a single isolate was collected and analyzed at each time point. Multiple isolates from each time point would provide more statistical confidence to our findings and could give more evidence for evolution within the patient. In conclusion, we identified intra-patient genetic variation of Mav during persistent human infection, and observed surprisingly high mutation rates. In addition, the response of the immune system to these clinical isolates was tested in macrophages and mice, demonstrating a highly adaptive microbe within the individual patients over time. Further investigation of the correlation between SNPs and adaptation of Mav may provide insight into the multiple strategies which Mav employs to resist chemotherapy and thereby help to develop more effective treatment strategies.

## MATERIAL AND METHODS

### Clinical isolates

40 clinical isolates from 15 patients diagnosed with MAC infections (2 to 6 consecutive isolates for each patient) were obtained from the Department of Medical Microbiology at St. Olavs Hospital in Trondheim, Norway. All isolates were previously characterized as MAC by the Norwegian Institute of Public Health in Oslo, Norway. The Regional Committee for Medical and Health Research Ethics approved this study (REK nord 2013/802).

The sixteen Mav isolates investigated in depth in the current study were collected in chronological order (2005-2007) from the sputum of patients. The isolates were grown on 7H10 Middlebrook (Difco/Becton Dickinson) medium supplemented with 10% ADC (Difco/Becton Dickinson). Single colonies were transferred to 7H9 Middlebrook (Difco/Becton Dickinson) liquid medium supplemented with 10% ADC (Difco/Becton Dickinson). All cultures were grown to the logarithmic phase OD_600nm_ 0.5-0.6; bacterial cultures were pelleted down and resuspended in PBS, followed by sonication, and finally passed through a syringe to obtain a single cell suspension. These suspensions were further used either in *in vitro* or *in vivo* infections.

### Pulsed Field Gel Electrophoresis (PFGE)

A modified protocol by Stevenson *et al*. (57) was employed. PFGE patterns were examined both visually and by computer-assisted analysis. Cluster analysis to compare SnaBI profiles and to construct dendrograms was performed using the Dice similarity coefficient and Unweighted Pair Group Method with Arithmetic Mean (UPGMA) in BioNumerics version 6.6 (Applied Maths, Sint-Martens-Latem, Belgium). General guidelines for interpreting chromosomal DNA restriction patterns were used in evaluating the relatedness of clinical isolates (58).

### Genome Sequencing and Assembly

DNA was extracted by the CTAB-lysozyme method (59). Samples were sequenced on an Illumina HiSeq 2500 with a read length of 106 bp or an Illumina HiSeq 4000 with a read length of 150 bp, both in paired-end mode. The mean depth of coverage was 109.0x (range 51-158). Genome sequences were assembled by a comparative-assembly method (60). Reads were mapped to Mav 104 (NC_008595.1) as a reference genome using BWA (61). Then, regions with indels or clusters of single nucleotide polymorphisms (SNPs) were identified and repaired by building local contigs from overlapping reads spanning such regions.

The sequence data for the clinical isolates has been uploaded to the NCBI Sequence Read Archive under accession number SRP136774 and are available through the following links https://www.ncbi.nlm.nih.gov/Traces/study/?acc=SRP136774,https://www.ncbi.nlm.nih.gov/bioproject/?term=PRJNA447977
,https://www.ncbi.nlm.nih.gov/sra?term=SRP136774

### Macrophages culture and infection

Bone marrow derived macrophages (BMDMs) were generated by culturing bone-marrow cells of C57BL/6 mice in RPMI-1640 medium (Sigma)/10% fetal calf serum (Gibco) and 20% L929 cell line supernatant for 4 days. Macrophages were seeded at 50,000 cells/ well in a 96 well plate and infected with Mav clinical isolates at a MOI of 10 for two hours. Cells were washed with Hanks balanced salt solution to eliminate extracellular bacteria. Three wells were lysed with PBS containing 0.02% Triton X (Sigma) and plated in serial dilutions on 7H10 Middlebrook plates in triplicate to enumerate uptake by CFU counting (t =0). At day 1, 3 and 7, infected cells were lysed and serial dilutions plated on 7H10 Middlebrook plates in triplicate, to record the course of infection.

### Mouse infection experiments

All protocols on animal work were approved by the Norwegian National Animal Research Authorities and carried out in accordance with Norwegian and European regulations and guidelines. C57BL/6 mice were bred in house and used at 6-8 weeks of age for experiments. Infection was performed by intraperitoneal injection of log-phase mycobacteria (2×10^7^−1×10^9^ CFU/mouse) in 0.2 ml PBS; inoculum was measured by CFU plating. At given time-points after infection, mice were killed, and spleen and liver were collected. Bacterial load was measured by plating serial dilutions of organ homogenates (spleen, liver) on 7H10 Middlebrook plates.

### Mav-specific T cell cytokine production

Splenocytes were isolated and prepared for flow cytometry as described in (33). Splenocytes were stimulated overnight with clinical Mav isolates from patient 9 and 13 at a MOI of 1. Concanavallin A (Sigma, 2.5µg/ml) stimulation was used as positive control, unstimulated cells served as negative control. Protein transport inhibitor (eBioscience) was added during the last 4h of stimulation. Surface antigen were stained with monoclonal antibodies against CD3 (FITC), CD4 (Brilliant Violet 605) and CD8 (Brilliant Violet 785, all from Biolegend). After fixation (2% PFA, Sigma) and permeabilization (0.5 %, Sigma), intracellular cytokine staining was performed for IFNγ (PE) and TNFα (APC, both from Biolegend). Flow-cytometry was performed on a BD LSR II flow-cytometer (BD Biosciences) and data analyzed using FlowJo (FlowJo, LLC) and GraphPad Prism (GraphPad Software, Inc.) software.

### Histopathology

Organ samples were fixed in buffered formalin, processed through standard dehydration, clearing and placed in paraffin overnight, cut in 5 µm thick sections and stained with hematoxylin & eosin. Microscopic images were taken with a Lumenera Infinity 2 camera and INFINITY ANALYZE software, release 6.2 (Lumenera Corp.) using a Nikon eclipse Ci microscope (Nikon) with 40x magnification.

## Statistics

All values from intracellular replication experiments are means of four independent experiments, performed in duplicate or triplicate. Results are presented as mean ± SEM and analyzed by two-way ANOVA followed by a Turkey post-test. Values obtained in mice experiments show mean± SEM of two independent experiments, with four mice in each group. Results were analyzed by two-way ANOVA, followed by Bonferroni post-test. In all analyses, a p-value of <0.05 was considered as statistically significant. Data analysis and statistical tests were performed with Graph Pad Prism 5.0 (GraphPad Software, Inc.).

## ACKNOWLEDGEMENTS

This work was partly supported by the Research Council of Norway through its Centres of Excellence funding scheme, project number 223255/F50. In addition the Research Council of Norway supported the work through project numbers 220836/F10 and 246944/F10. In addition NTNU and the Liaison Committee between the Central Norway Regional Health Authority (RHA) and the Norwegian University of Science and Technology (NTNU) contributed with funding to the study.

## Supplementary Figure Legends

**FIG S1** Differential melting curves from 16S rRNA gene qRT-PCR. The curves of 19 clinical MAC isolates as well as positive (*M. intracellulare* and Mtb) and negative controls are shown.

**FIG S2** Bone marrow derived murine macrophages were infected with clinical stains, collected in chronological order, at a MOI of 10. After two hours of infection, cells were washed and lysed to analyze uptake. Uptake is represented as log CFU and designated as zero hour.

**FIG S3** Nanostring data. Murine BMDMs were infected with the first and last clinical isolate of Mav (9.1 and 9.6) at a MOI of 10. After 6 hours post infection, cells were lysed in RLT buffer and subsequently RNA was extracted to perform nanostring analysis. nCounter® GX Mouse Inflammation Kit was used and the results were analyzed on *Partek*^®^ *Genomics Suite*^®^ software.

**FIG S4** Proinflammatory cytokines IL-6 (A) and IL-1β (B) measured from liver homogenates. Results are presented as ng cytokine per gram tissue.

**FIG S5** Accumulated number of SNPs, accounting for sub clones. SNPs were counted relative to Mav 104 for patients 9 and 13. The isolates from patient 13 were treated as two separate lineages (13.1-2 and 13.3-4). For patient 4, SNPs in isolates 4.2-4.6 were counted relative to 4.2.

**Table S1** Medical history and time line of sample collection for all the patients in the study.

**Table S2** List of genes containing SNPs. Both synonymous and non-synonymous mutations were observed. Putative gene function mentioned for non-synonymous mutations. Mav 104 reference from KEGG database was used to annotate function.

**Table S3** List of genes containing mutations (relative to Mav104, as an outgroup). Both, synonymous and non-synonymous mutations were observed. Putative gene function mentioned for non-synonymous mutations. Mav 104 reference from KEGG database was used to annotate function. The mutations for patient 13 are highlighted in red and blue to emphasize the two sub-series of isolates, (13.1, and 13.2) and (13.3, and13.4). The mutations for patient 4 are highlighted in red and blue to emphasize the two sub-series of isolates (4.1) and (4.2-4.6)

## REFERENCES

1. Inderlied CB, Kemper CA, Bermudez LE. 1993. The Mycobacterium avium complex. Clin Microbiol Rev 6:266–310.

2. Falkinham JO. 2009. Surrounded by mycobacteria: Nontuberculous mycobacteria in the human environment. J Appl Microbiol 107:356–367.

3. Faria S, Joao I, Jordao L. 2015. General Overview on Nontuberculous Mycobacteria, Biofilms, and Human Infection. J Pathog 2015:1–10.

4. Horsburgh CR, Gettings J, Alexander LN, Lennox JL. 2001. Disseminated Mycobacterium avium Complex Disease among Patients Infected with Human Immunodeficiency Virus, 1985 – 2000 2118:3–8.

5. Tateishi Y, Hirayama Y, Ozeki Y, Nishiuchi Y, Yoshimura M, Kang J, Shibata A, Hirata K, Kitada S, Maekura R, Ogura H, Kobayashi K, Matsumoto S. 2009. Virulence of Mycobacterium avium complex strains isolated from immunocompetent patients. Microb Pathog 46:6–12.

6. Field SK, Fisher D, Cowie RL. 2004. Mycobacterium avium complex Pulmonary Disease in Patients Without HIV Infection. Chest 126:566–581.

7. Bruijnesteijn Van Coppenraet LES, De Haas PEW, Lindeboom JA, Kuijper EJ, Van Soolingen D. 2008. Lymphadenitis in children is caused by Mycobacterium avium hominissuis and not related to “bird tuberculosis.” Eur J Clin Microbiol Infect Dis 27:293–299.

8. Griffith DE, Aksamit T, Brown-Elliott BA, Catanzaro A, Daley C, Gordin F, Holland SM, Horsburgh R, Huitt G, Iademarco MF, Iseman M, Olivier K, Ruoss S, Von Reyn CF, Wallace RJ, Winthrop K. 2007. An official ATS/IDSA statement: Diagnosis, treatment, and prevention of nontuberculous mycobacterial diseases. Am J Respir Crit Care Med 175:367–416.

9. Ryu YJ, Koh WJ, Daley CL. 2016. Diagnosis and treatment of nontuberculous mycobacterial lung disease: Clinicians’ perspectives. Tuberc Respir Dis (Seoul) 79:74–84.

10. Brode SK, Daley CL, Marras TK. 2014. The epidemiologic relationship between tuberculosis and non-tuberculous mycobacterial disease: a systematic review. Int J Tuberc Lung Dis 18:1370–1377.

11. Cooper AM, Mayer-Barber KD, Sher A. 2011. Role of innate cytokines in mycobacterial infection. Mucosal Immunol 4:252–260.

12. Furney SK, Skinner PS, Roberts AD, Appelberg R, Orme IM. 1992. Capacity of Mycobacterium avium isolates to grow well or poorly in murine macrophages resides in their ability to induce secretion of tumor necrosis factor. Infect Immun.

13. Appelberg R, Castro a. G, Pedrosa J, Silva R a., Orme IM, Minoprio P. 1995. Erratum: Role of gamma interferon and tumor necrosis factor alpha during T-cell-independent and -dependent phases of Mycobacterium avium infection (Infection and Immunity 62:9 (3969)). Infect Immun 63:1145.

14. Awuh JA, Flo TH. 2017. Molecular basis of mycobacterial survival in macrophages. Cell Mol Life Sci 74:1625–1648.

15. Mayer-Barber KD, Sher A. 2015. Cytokine and lipid mediator networks in tuberculosis. Immunol Rev 264:264–275.

16. Appelberg R. 1994. Protective Role of Interferon Gamma, Tumor Necrosis Factor Alpha and Interleukin-6 in Mycobacterium tuberculosis and M. avium Infections. Immunobiology 191:520–525.

17. Van Heyningen TK, Collins HL, Russell DG. 1997. IL-6 Produced by Macrophages Infected with Mycobacterium species Suppresses T Cell Responses. J Immunol 158:330–337.

18. Fattorini L, Xiao Y, Li B, Santoro C, Ippoliti F, Orefici G. 1994. Induction of IL-1??, IL-6, TNF-??, GM-CSF and G-CSF in human macrophages by smooth transparent and smooth opaque colonial variants of Mycobacterium avium. J Med Microbiol 40:129–133.

19. Ernst JD, Trevejo-nuñez G, Banaiee N. 2007. Science in medicine Genomics and the evolution, pathogenesis, and diagnosis of tuberculosis. J Clin Invest 117:1738–1745.

20. Ford CB, Lin PL, Chase MR, Shah RR, Iartchouk O, Galagan J, Mohaideen N, Ioerger TR, Sacchettini JC, Lipsitch M, Flynn JL, Fortune SM. 2011. Use of whole genome sequencing to estimate the mutation rate of Mycobacterium tuberculosis during latent infection. Nat Genet 43:482–486.

21. Uchiya K, Tomida S, Nakagawa T, Asahi S, Nikai T, Ogawa K. 2017. Comparative genome analyses of Mycobacterium avium reveal genomic features of its subspecies and strains that cause progression of pulmonary disease. Sci Rep 7:39750.

22. Semret M, Zhai G, Mostowy S, Cleto C, Alexander D, Cangelosi G, Cousins D, Collins DM, Van Soolingen D, Behr MA. 2004. Extensive Genomic Polymorphism within Mycobacterium avium. J Bacteriol 186:6332–6334.

23. Oliveira RS, Sircili MP, Oliveira EMD, Balian SC, Ferreira-Neto JS, Leão SC. 2003. Identification of Mycobacterium avium Genotypes with Distinctive Traits by Combination of IS1245-Based Restriction Fragment Length Polymorphism and Restriction Analysis of hsp65. J Clin Microbiol 41:44–49.

24. Pedrosa J, Flórido M, Kunze ZM, Castro a G, Portaels F, McFadden J, Silva MT, Appelberg R. 1994. Characterization of the virulence of Mycobacterium avium complex (MAC) isolates in mice. Clin Exp Immunol 98:210–6.

25. Amaral EP, Kipnis TL, de Carvalho ECQ, da Silva WD, Leão SC, Lasunskaia EB. 2011. Difference in virulence of mycobacterium avium isolates sharing indistinguishable dna fingerprint determined in murine model of lung infection. PLoS One 6.

26. Zhao X, Epperson LE HN, Honda JR, Chan ED, Strong M WN, RM. D. 2017. Complete Genome Sequence of Mycobacterium aviumsubsp.hominissuis Strain H87 Isolated from an Indoor Water Sample. Genome Announc 5:16–17.

27. Walker TM, Ip CLC, Harrell RH, Evans JT, Kapatai G, Dedicoat MJ, Eyre DW, Wilson DJ, Hawkey PM, Crook DW, Parkhill J, Harris D, Walker a. S, Bowden R, Monk P, Smith EG, Peto TE a. 2013. Whole-genome sequencing to delineate Mycobacterium tuberculosis outbreaks: A retrospective observational study. Lancet Infect Dis 13:137–146.

28. Appelberg R. 2006. Pathogenesis of Mycobacterium avium infection: typical responses to an atypical mycobacterium? Immunol Res 35:179–190.

29. Awuh JA, Haug M, Mildenberger J, Marstad A, Do CPN, Louet C, Stenvik J, Steigedal M, Damås JK, Halaas Ø, Flo TH. 2015. Keap1 regulates inflammatory signaling in *Mycobacterium avium* -infected human macrophages. Proc Natl Acad Sci 112:E4272–E4280.

30. Gidon A, Åsberg SE, Louet C, Ryan L, Haug M, Flo TH. 2017. Persistent mycobacteria evade an antibacterial program mediated by phagolysosomal TLR7/8/MyD88 in human primary macrophages. PLoS Pathog 13:1–27.

31. Blumenthal A, Lauber J, Hoffmann R, Ernst M, Keller C, Buer J, Ehlers S, Reiling N. 2005. Common and unique gene expression signatures of human macrophages in response to four strains of Mycobacterium avium that differ in their growth and persistence characteristics. Infect Immun.

32. Doherty TM, Sher a. 1997. Defects in cell-mediated immunity affect chronic, but not innate, resistance of mice to Mycobacterium avium infection. J Immunol 158:4822–4831.

33. Haug M, Awuh J a., Steigedal M, Frengen Kojen J, Marstad A, Nordrum IS, Halaas Ø, Flo TH. 2013. Dynamics of immune effector mechanisms during infection with mycobacterium avium in C57BL/6 mice. Immunology 140:232–243.

34. Ichikawa K, Yagi T, Moriyama M, Inagaki T, Nakagawa T, Uchiya KI, Nikai T, Ogawa K. 2009. Characterization of Mycobacterium avium clinical isolates in Japan using subspecies-specific insertion sequences, and identification of a new insertion sequence, ISMav6. J Med Microbiol 58:945–950.

35. Ichikawa K, van Ingen J, Koh WJ, Wagner D, Salfinger M, Inagaki T, Uchiya K ichi, Nakagawa T, Ogawa K, Yamada K, Yagi T. 2015. Genetic diversity of clinical Mycobacterium avium subsp. hominissuis and Mycobacterium intracellulare isolates causing pulmonary diseases recovered from different geographical regions. Infect Genet Evol 36:250–255.

36. Uchiya K ichi, Takahashi H, Yagi T, Moriyama M, Inagaki T, Ichikawa K, Nakagawa T, Nikai T, Ogawa K. 2013. Comparative Genome Analysis of Mycobacterium avium Revealed Genetic Diversity in Strains that Cause Pulmonary and Disseminated Disease. PLoS One 8.

37. Ford CB, Shah RR, Maeda MK, Gagneux S, Murray MB, Cohen T, Johnston JC, Gardy J, Lipsitch M, Fortune SM. 2013. Mycobacterium tuberculosis mutation rate estimates from different lineages predict substantial differences in the emergence of drug-resistant tuberculosis. Nat Genet 45:784–790.

38. Iwamoto T, Nakajima C, Nishiuchi Y, Kato T, Yoshida S, Nakanishi N, Tamaru A, Tamura Y, Suzuki Y, Nasu M. 2012. Infection, Genetics and Evolution Genetic diversity of Mycobacterium avium subsp. hominissuis strains isolated from humans, pigs, and human living environment. Infect Genet Evol 12:846–852.

39. Pérez-Lago L, Comas I, Navarro Y, González-Candelas F, Herranz M, Bouza E, García-De-Viedma D. 2014. Whole Genome Sequencing Analysis of Intrapatient Microevolution in Mycobacterium tuberculosis: Potential Impact on the Inference of Tuberculosis Transmission. J Infect Dis.

40. Liu Q, Via LE, Luo T, Liang L, Liu X, Wu S, Shen Q, Wei W, Ruan X, Yuan X, Zhang G, Barry Iii CE, Gao Q. 2015. Within patient microevolution of Mycobacterium tuberculosis correlates with heterogeneous responses to treatment. Nat Publ Gr.

41. Eldholm V, Norheim G, von der Lippe B, Kinander W, Dahle UR, Caugant DA, Mannsåker T, Mengshoel AT, Dyrhol-Riise AM, Balloux F. 2014. Evolution of extensively drug-resistant Mycobacterium tuberculosisfrom a susceptible ancestor in a single patient. Genome Biol 15:490.

42. O’Neill MB, Mortimer TD, Pepperell CS. 2015. Diversity of Mycobacterium tuberculosis across Evolutionary Scales. PLoS Pathog 11:1–29.

43. Crowle AJ, Tsang AY, Vatter AE, May MH. 1986. Comparison of 15 laboratory and patient-derived strains of Mycobacterium avium for ability to infect and multiply in cultured human macrophages. J Clin Microbiol.

44. Birkness KA, Swords WE, Huang PH, White EH, Dezzutti CS, Lal RB, Quinn FD. 1999. Observed differences in virulence-associated phenotypes between a human clinical isolate and a veterinary isolate of Mycobacterium avium. Infect Immun 67:4895–4901.

45. Tse HM, Josephy SI, Chan ED, Fouts D, Cooper AM. 2002. Activation of the mitogen-activated protein kinase signaling pathway is instrumental in determining the ability of Mycobacterium avium to grow in murine macrophages. J Immunol 168:825–833.

46. Petrofsky M, Bermudez LE. 1999. Neutrophils fromMycobacterium avium-Infected Mice Produce TN F-α, IL-12, and IL-1β and Have a Putative Role in Early Host Response. Clin Immunol 91:354–358.

47. Roque S, Nobrega C, Appelberg R, Correia-Neves M. 2007. IL-10 underlies distinct susceptibility of BALB/c and C57BL/6 mice to Mycobacterium avium infection and influences efficacy of antibiotic therapy. J Immunol 178:8028–8035.

48. Flórido M, Correia-Neves M, Cooper AM, Appelberg R. 2003. The cytolytic activity of natural killer cells is not involved in the restriction of Mycobacterium avium growth. Int Immunol 15:895–901.

49. Pellegrin JL, Taupin JL, Dupon M, Ragnaud JM, Maugein J, Bonneville M, Moreau JF. 1999. Gammadelta T cells increase with Mycobacterium avium complex infection but not with tuberculosis in AIDS patients. Int Immunol 11:1475–1478.

50. Lockhart E, Green AM, Flynn JL. 2006. IL-17 production is dominated by gammadelta T cells rather than CD4 T cells during Mycobacterium tuberculosis infection. J Immunol 177:4662–9.

51. Gilbertson B, Zhong J, Cheers C. 1999. Anergy, IFN-gamma production, and apoptosis in terminal infection of mice with Mycobacterium avium. J Immunol 163:2073–80.

52. Zhong J, Gilbertson B, Cheers C. 2003. Apoptosis of CD4+ and CD8+ T cells during experimental infection with Mycobacterium avium is controlled by Fas/FasL and Bcl-2-sensitive pathways, respectively. Immunol Cell Biol 81:480–486.

53. Horwitz MA, Lee BW, Dillon BJ, Harth G. 1995. Protective immunity against tuberculosis induced by vaccination with major extracellular proteins of Mycobacterium tuberculosis. Proc Natl Acad Sci 92:1530–1534.

54. Liu X, Feng Z, Harris NB, Cirillo JD, Bercovier H, Barletta RG. 2001. Identification of a secreted superoxide dismutase in Mycobacterium avium ssp. paratuberculosis. FEMS Microbiol Lett 202:233–238.

55. Sherman DR, Sabo PJ, Hickey MJ, Arain TM, Mahairas GG, Yuant Y, Barry, 3rd CE, Stover CK. 1995. Disparate responses to oxidative stress in saprophytic and pathogenic mycobacteria. Proc Natl Acad Sci U S A 92:6625–6629.

56. Geier H, Mostowy S, Cangelosi GA, Behr MA, Ford TE. 2008. Autoinducer-2 triggers the oxidative stress response in Mycobacterium avium, leading to biofilm formation. Appl Environ Microbiol 74:1798–1804.

57. Stevenson K, Hughes VM, Juan L De, Inglis NF, Wright F, Sharp JM. 2002. Molecular Characterization of Pigmented and Nonpigmented Isolates of Mycobacterium avium subsp 1. paratuberculosis. Society 40:1798–1804.

58. Tenover FC, Arbeit RD, Goering R V., Mickelsen PA, Murray BE, Persing DH, Swaminathan B. 1995. Interpreting chromosomal DNA restriction patterns produced by pulsed-field gel electrophoresis: Criteria for bacterial strain typing. J Clin Microbiol 33:2233–2239.

59. Larsen MH, Biermann K, Tandberg S, Hsu T, Jacobs WR. 2007. Genetic Manipulation of Mycobacterium tuberculosis. Curr Protoc Microbiol.

60. Ioerger TR, Feng Y, Ganesula K, Chen X, Dobos KM, Fortune S, Jacobs WR, Mizrahi V, Parish T, Rubin E, Sassetti C, Sacchettini JC. 2010. Variation among genome sequences of H37Rv strains of Mycobacterium tuberculosis from multiple laboratories. J Bacteriol 192:3645–3653.

61. Li H, Durbin R. 2009. Fast and accurate short read alignment with Burrows-Wheeler transform. Bioinformatics 25:1754–1760.

